# Neuropsychological assessment and virtual reality training of social prediction in patients with cerebellar malformation

**DOI:** 10.1101/2021.01.19.427247

**Authors:** Cosimo Urgesi, Niccolò Butti, Alessandra Finisguerra, Emilia Biffi, Enza Maria Valente, Romina Romaniello, Renato Borgatti

## Abstract

It has been proposed that impairments of the predictive function exerted by the cerebellum may account for social cognition deficits. Here, we integrated cerebellar functions in a predictive coding framework to elucidate how cerebellar alterations could affect the predictive processing of others’ behavior. Experiment 1 demonstrated that cerebellar patients were impaired in relying on contextual information during action prediction, and this impairment was significantly associated with social cognition abilities. Experiment 2 indicated that patients with cerebellar malformation showed a domain-general deficit in using contextual information to predict both social and physical events. Experiment 3 provided first evidence that a social-prediction training in virtual reality could boost the ability to use context-based predictions to understand others’ intentions. These findings shed new light on the predictive role of the cerebellum and its contribution to social cognition, paving the way for new approaches to the rehabilitation of the Cerebellar Cognitive Affective Syndrome.

## Introduction

The Cerebellar Cognitive Affective Syndrome (CCAS; (Schmahmann & Sherman, 1998)), refers to a complex constellation of deficits in executive functions, visuo-spatial skills, language, affect and behavior regulation, all characterized by an augmented or diminished response to external and internal stimuli (Schmahmann, 2004). CCAS symptoms (Schmahmann et al., 2007) include deficits in social behavior and at diverse levels of social cognitive processing, such as action perception (Abdelgabar et al., 2019; Cattaneo et al., 2012), emotion recognition (Adamaszek et al., 2017; D’Agata et al., 2011; Hoche et al., 2016) and theory of mind (ToM) abilities (Clausi et al., 2019; Sokolovsky et al., 2010).

To explain how cerebellar alterations could result in this wide variety of cognitive, linguistic and social symptoms, Schmahmann (1996, 2004) proposed that the cerebellum plays a domaingeneral, supramodal computation defined as Universal Cerebellar Transform (UCT). This hypothesis was grounded on the classical models of the cerebellum in sensorimotor control (Albus, 1971; M. Ito, 1984; Marr, 1969), which postulated that the cerebellum could act as a general regulator of motor activity through its primary functions of predictive internal models and error-signaling (Miall & Wolpert, 1996). In this view, the cerebellum plays a general role in maintaining behavior around a homeostatic baseline according to context, Although the UCT hypothesis has been questioned for its account of cerebellar functionality (Diedrichsen et al., 2019), it has provided an innovative framework to understand the complex pattern of motor, cognitive and social deficits following cerebellar alterations (Manto & Mariën, 2015; Schmahmann, 2019; Sokolov et al., 2017).

Predictive coding accounts postulate that the brain constantly generates predictions about incoming events, working as a Bayesian inference machine (Knill & Pouget, 2004). Through reciprocal and recurrent interactions at any stage of stimulus processing, bottom-up sensorial information is compared with top-down expectations, the so-called priors, which arise from innate assumptions (e.g., the light come from above) or from implicitly learned contextual regularities (Series & Seitz, 2013). Mismatches between current sensorial evidence and priors produce predictionerror signals that inform subsequent layers and update top-down predictions about the expected input, until prediction error is minimized (Friston, 2012b). Thus, priors offer contextual guidance to select the most probable cause for that specific sensorial input (Kilner et al., 2007b). This theoretical framework has demonstrated its reliability in accounting for perception (Friston & Kiebel, 2009), interoception (Seth & Friston, 2016), and sensorimotor control (Körding & Wolpert, 2004) as well as social cognition (Brown & Brüne, 2012).

Indeed, social interactions continuously require individuals to anticipate others’ actions according to context and past experience, and to predict others’ intentions from ambiguous movement kinematics (Amoruso & Finisguerra, 2019). Noteworthy, predictive coding provides a unique platform to understand social processing at diverse levels of complexity, from action perception (Friston et al., 2011) to ToM (Koster-Hale & Saxe, 2013). Predictive coding accounts of the Action Observation Network (AON; (Friston et al., 2011; Kilner et al., 2007a; Neal & Kilner, 2010)) propose that we understand others’ actions by inverting forward models of action execution, since actions driven with different intentions are performed with different movement kinematics (Catmur, 2015; Koul et al., 2018). In this sense, the activation of fronto-parietal corticomotor areas during action observation (Rizzolatti & Craighero, 2004) and the deficits of patients with fronto-parietal lesions in action understanding (Avenanti et al., 2013) would point to the importance of motor simulation in action perception.

Research on cerebellar patients has reported deficits in action understanding (Abdelgabar et al., 2019; Cattaneo et al., 2012; Sokolov et al., 2010), sustaining that the cerebellum is directly involved in the simulation of biological movement, particularly when kinematics provides subtle clues. However, the impairment of cerebellar patients in action perception could be specifically explained by an impairment of predictive processing (Sokolov, 2018). Indeed, the cerebellum might integrate contextual and movement information into a-priori predictive models that modulate cognitive processing of others’ actions, allowing to overcome kinematics uncertainty Crucially, if an impairment of context-based predictions following cerebellar alterations results in limited ability to understand others’ actions, emotions, and mental states (Butti, Corti, et al., 2020), rehabilitative approaches may target this function to treat the behavioral disorders and social cognition deficits encompassed by the CCAS (Schmahmann, 2010).

Considering these premises, our work aimed to investigate whether cerebellar alterations could impair context-based prediction of social events (Experiment 1) and whether this deficit was differently reliable for social and physical events (Experiment 2). Thus, we aimed to examine the efficacy of a rehabilitative intervention in virtual reality (VR) targeting social prediction (Butti, Biffi, et al., 2020) in boosting the use of contextual priors during action prediction in cerebellar patients (Experiment 3).

## Results

### Experiment 1: Action prediction task and social cognition

Experiment 1 was aimed at testing the hypothesis that cerebellar alterations affect the integration of contextual and kinematics information into context-based predictions (i.e., priors) of others’ actions. To this aim, we administered to cerebellar pediatric patients (CM; N =26), to age- and cognitive-level-matched patients with congenital neurological disorders not affecting the cerebellum (CND; N=26), and to healthy peers (TD; N=26), an action prediction task already adopted in evaluating the use of contextual priors in children with autism (Amoruso et al., 2019). Demographic and clinical variables of the tree group are reported in Table 1.

**Table 1.**
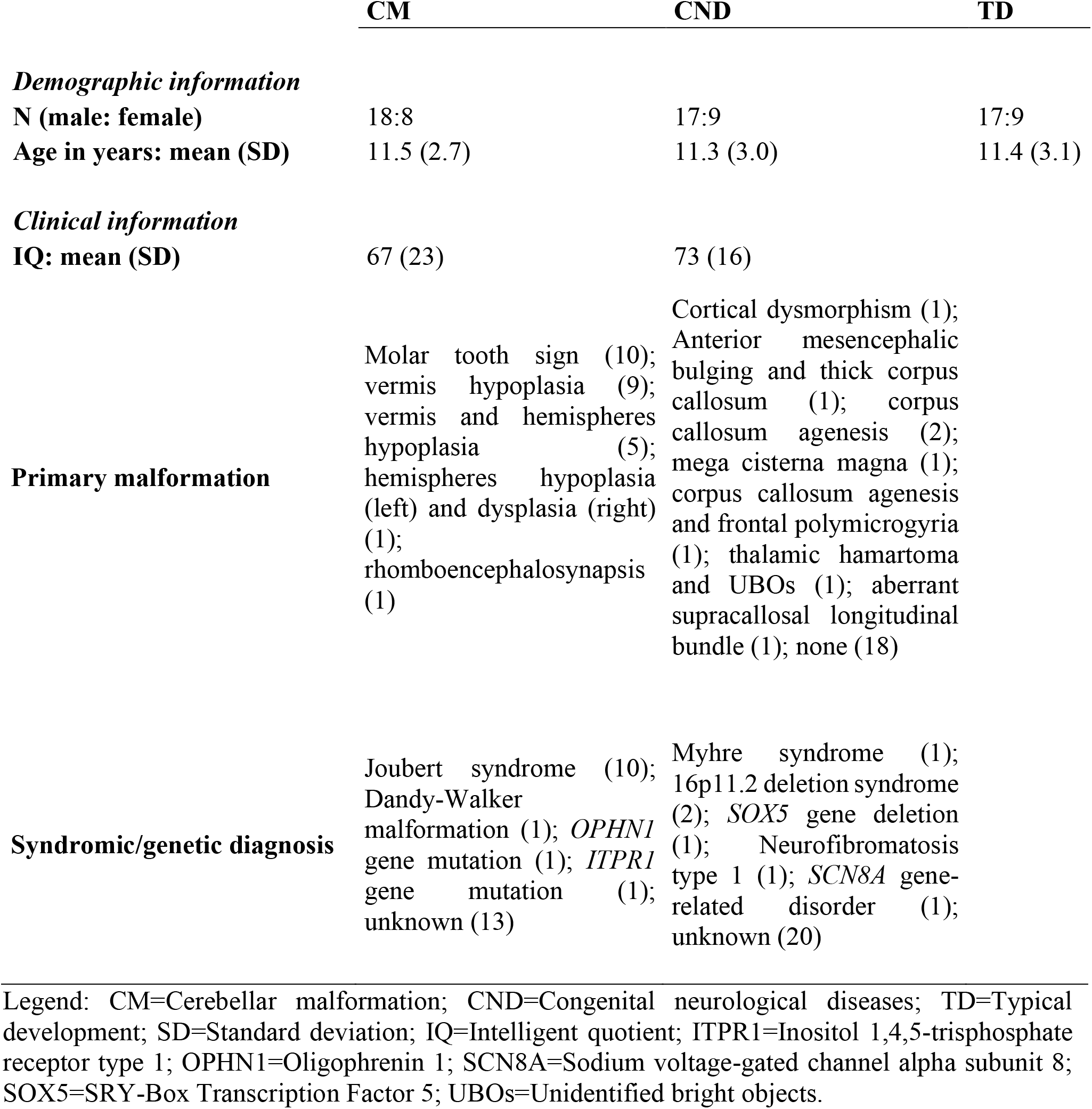
Demographic and clinical information of the three groups in experiment 1.

Briefly, the task uses an implicit learning procedure (familiarization phase) to allow participants to learn statistical regularities of co-occurrence (i.e., 10%, 40%, 60%, 90%) between action intentions and contextual cues and then tests (testing phase) the use of these contextual priors in condition of perceptual ambiguity (see Fig. 1a,b and Material and Methods for detailed information). Furthermore, we administered patients with the NEPSY-II social perception subtests assessing verbal and non-verbal ToM abilities and facial affect recognition (Korkman et al., 2007). We expected that only cerebellar patients should show a deficit in relying on contextual priors during action perception. Furthermore, the use of context-based predictions for social stimuli should be associated with non-verbal social perception abilities, leading to lower performances of cerebellar participants than control patients and healthy peers.

**Figure 1.**
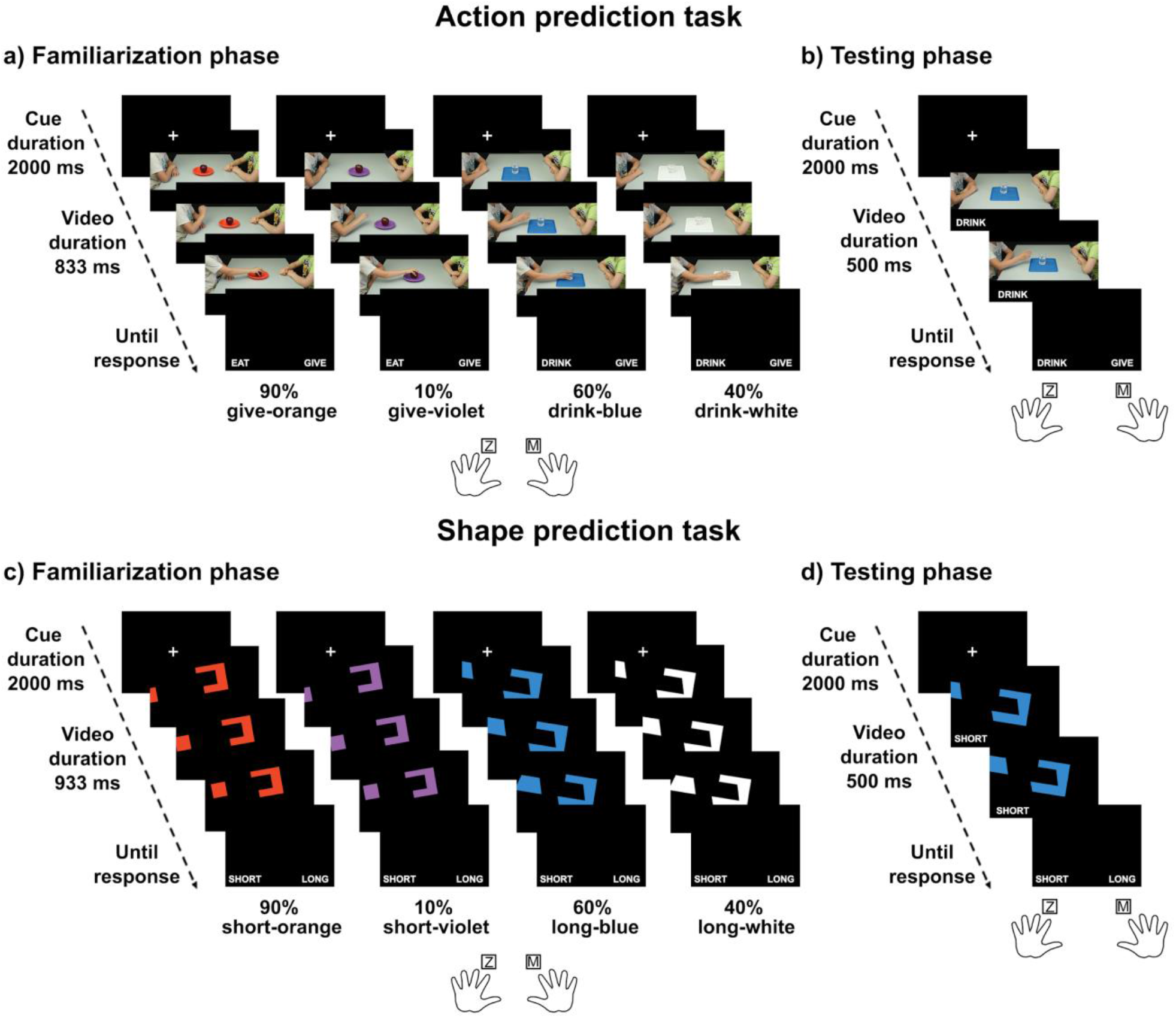
Structure of the predictive tasks. Trial structure, timeline and examples of probabilistic contextual cue-action associations in the familiarization and testing phase of the action prediction task (a, b) and the shape prediction task (c, d).

Preliminary analyses confirmed that groups did not differ for age (p=0.961) and gender (p=0.944). For the familiarization phase, the one-way, between-subjects ANOVA revealed significant effect of group (F_2,75_=7.13, p=0.001, η^2^_p_=0.16), with CM patients (90.62±1.63%) less accurate than CND (95.73±0.80%; p=0.003) and TD participants (96.16±0.84%; p=0.002), but without differences between the two control groups (p=0.796). While these findings confirmed that alterations of the cerebellum affect perception of biological movements (Abdelgabar et al., 2019; Cattaneo et al., 2012; Ferrari et al., 2019), accuracy of the CM participants remained high (>90%), ensuring that they were exposed to the probabilistic action-context associations. Crucially, in the testing phase, only CM patients failed to show a contextual modulation on their responses, even if the amount of kinematic information was exactly the same across probability conditions. Indeed, the mixed-model ANOVA with group as between-subject factor and probability (10 vs. 40 vs. 60 vs. 90%) as within-subjects variable yielded significant effects of group (F_2,75_=8.47, p<0.001, η^2^_p_=0.18) and probability (F_3,225_=6.95, p<0.001, η^2^_p_=0.08), better qualified by their interaction (F_6, 225_=4.36, p<0.001, η^2^_p_=0.1). TD participants showed significantly higher accuracy for the 90% action-context co-occurrence (86.00±2.10%) compared to the 10% one (71.23±3.21%; p<0.011). This latter condition was also lower than both the 40% (81.08±2.97%; p=0.011) and 60% (78.87±3.53%; p=0.040) conditions. Similarly, CND patients were less accurate in the 10% condition (66.00±4.68%) compared to the 90% (83.12±2.62%; p<0.001), 40% (79.27±3.52%; p<0.001) and 60% (74.65±4.30%; p=0.024) ones. Furthermore, their accuracy was higher in the 90% than in the 60% condition (p=0.031). Conversely, no difference emerged within the CM group (10%: 68.08±3.69%, 40%: 62.81±3.41%, 60%: 63.31±3.27%, 90%: 63.42±3.25%; all p>0.190). Between-group comparisons showed that, in the lowest-probability condition, the three groups had comparable performance (all p>0.314), while for all the others probability associations CND and TD participants showed comparable accuracy (all p>0.386), but outperformed CM patients (all p<0.040). This rules out that the performance of CM patients can be uniquely explained by their difficulties in perceiving biological movements and validated our hypothesis that cerebellar alterations impair context-based predictions of social events (Fig. 2).

**Figure 2.**
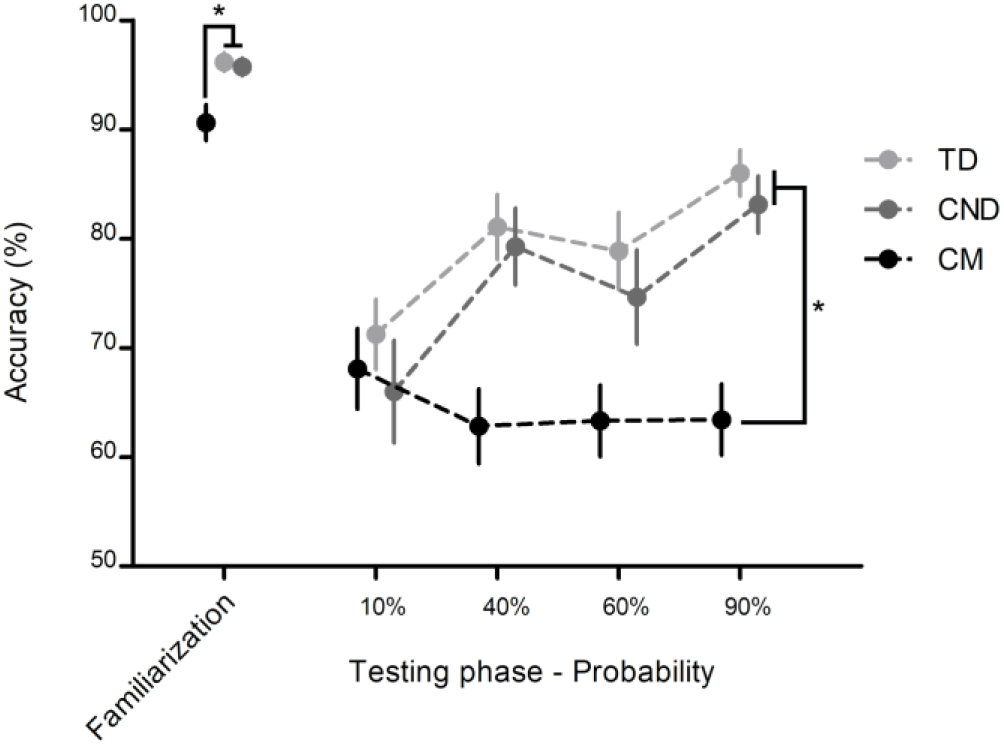
Results of the action prediction task in Experiment 1. Twenty-six patients per group were administered with the action prediction task. Trials with anticipated or out-of-time responses (RT<150 ms or >5,000 ms) were excluded from analysis. Accuracy values for the familiarization and testing phases were treated, respectively, with between-subjects and mixed model ANOVA designs. Asterisks indicate significant between-group comparisons for the familiarization (p=0.001) and for the highest-probability condition of the testing phase (p<0.001).

To correlate performance at the task and social perception abilities, in line with previous research (Amoruso et al., 2019; Butti, Corti, et al., 2020), we calculated a standardized beta coefficient across trials of the testing phase that represents the modulatory effect of the probabilistic associations, thus providing a measure of the strength of the contextual priors. Correlation analyses (Table 2) revealed that for CND patients the beta index was positively associated with *T*-scores in the nonverbal part of ToM subtest (r=0.46, p=0.026) and in the affect recognition subtest (r=0.42, p=0.046). This supports the importance of effective predictive processing of social events also for higher-level social perception skills. It is to note that these relationships were not reliable in CM patients (all p>0.130), likely due to the flattened performance of this group in the action prediction task. Fisher’s Z-transformations confirmed that these correlations were significantly different between the two clinical groups (−2.48≤Z≤2.22; p<0.013) (Fig. 3).

**Figure 3.**
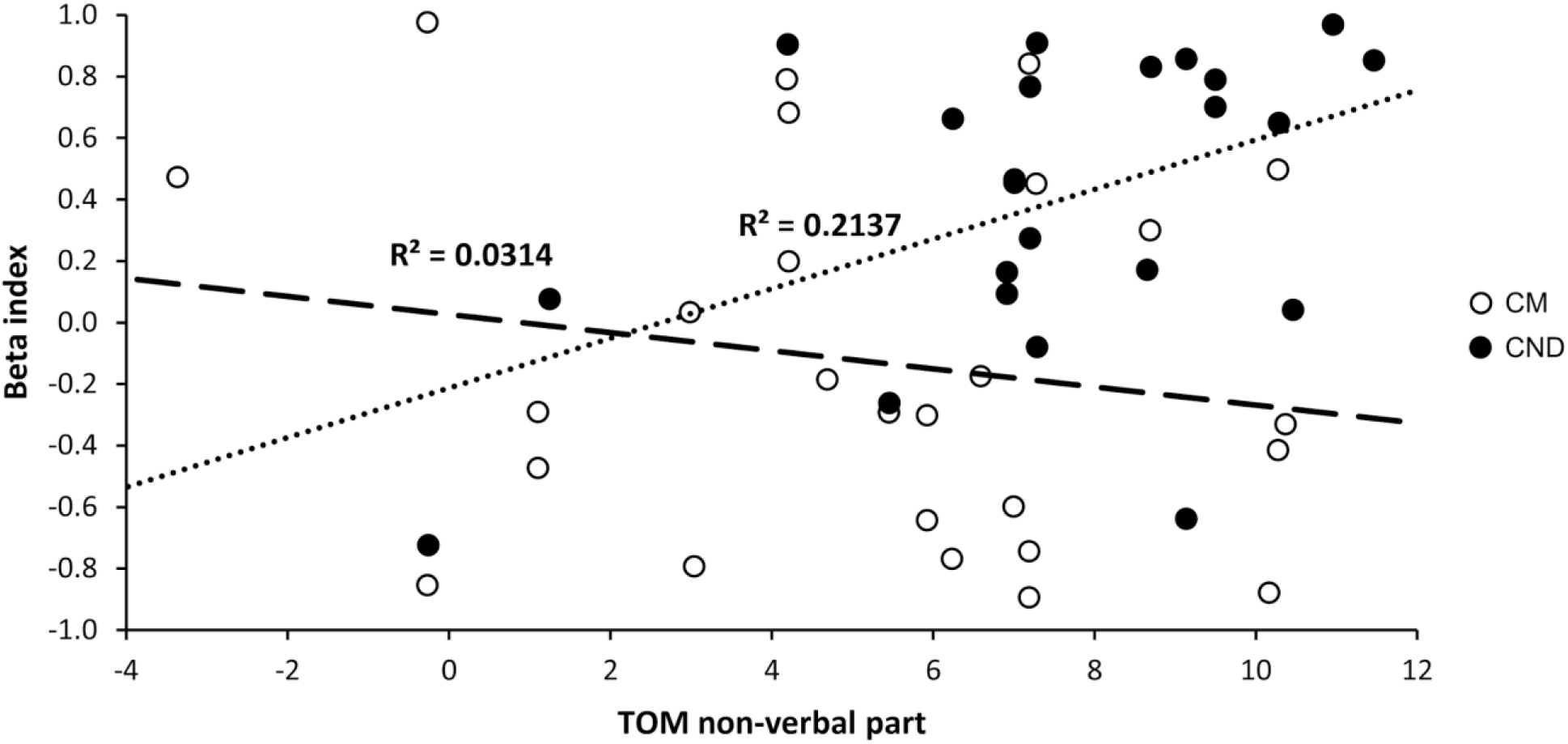
Correlations between beta index and non-verbal ToM abilities in cerebellar and control patients. The beta index was calculated across trials of the testing phase by running, at individual level, a regression analysis with probability as predictor and accuracy as dependent variable, thus representing a measure of the strength of the contextual priors. Please note that three CND patients were not administered with the ToM subtest (N=23; N of CM patients=26). Circles represent individual values; thicker dotted black line represents the correlation for CM patients (p=0.39), thinner dotted black line represents the correlation for CND patients (p=0.026).

**Table 2.** IQ, social perception abilities, and their correlations with the beta index in the two clinical groups. Significant results are reported in bold.

|  | CM |  | CND |  | Group Comparison |
| --- | --- | --- | --- | --- | --- |
|  | mean (SD) | Correlation with beta r (p) | mean (SD) | Correlation with beta r (p) | t<sub>50</sub> (p) |
| *General intellectual functioning (FSIQ)* | 67 (23) | -0.37 (0.06) | 73 (16) | 0.05 (0.84) | 1.01 (0.32) |
| *Social perception (t-scores)* |  |  |  |  |  |
| **ToM - verbal part** | 5.6 (5.0) | -0.06 (0.77) | 5.6 (5.2) | 0.18 (0.43) | 0.01 (0.99) |
| **ToM - non-verbal part** | 5.3 (3.6) | -0.18 (0.39) | 7.5 (2.9) | **0.46 (0.026)** | **2.45 (0.018)** |
| **Affect recognition** | 4.7 (4.3) | -0.30 (0.13) | 5.0 (5.1) | **0.42 (0.046)** | 0.26 (0.79) |

Legend: FSIQ=Full scale intelligent quotient; CM=Cerebellar malformations; CND=Congenital neurological disorders; SD=Standard deviation; ToM=Theory of Mind.

The clinical relevance of the prediction deficit in the social domain was confirmed by the performance in the non-verbal ToM subtest, in which CM patients were significantly impaired in inferring others’ mental states from contexts as compared to the control clinical group (t_50_=2.45, p=0.018, Cohen’s d=0.68), while all other comparisons were non-significant (all t<1.02, p>0.314). Moreover, no relationship was found between intellectual functioning and the beta index either overall or within the clinical groups, thus confirming a prediction deficit for CM patients regardless of their cognitive level.

Globally, results of Experiment 1 were in accordance with a predictive coding account of cerebellar contributions to social cognition (Butti, Corti, et al., 2020; Clausi et al., 2019). However, it was not clear whether this predictive deficit was specific for social stimuli or represented an impairment of a domain-general computation exerted by the cerebellum.

### Experiment 2: comparison of context-based predictions for social and physical events

In this experiment, we aimed to qualify the results of Experiment 1 by investigating whether the deficits of CM patients in using contextual priors are specific for action prediction or, alternatively, reflect the impairment of a domain-general predictive mechanism. We, thus, administered 18 CM patients and two control groups of 18 CND patients without cerebellar alterations and of 18 TD peers (see Table 3) with the same task as in Experiment 1 and with a shape prediction task developed to assess the use of contextual priors for predicting physical events (Bianco et al., 2020). The structure of the shape prediction task was similar to the action prediction task, but participants were required to predict the unfolding of moving geometrical shapes (see Fig. 1c,d and Material and Methods for detailed information). In line with the UCT hypothesis (Schmahmann, 1996, 2019), we expected that CM patients should fail to show a contextual modulation of their prediction of both social and physical events. However, previous research reported that cerebellar patients may present stronger difficulties in processing social stimuli than inanimate objects (Cattaneo et al., 2012; Sokolov et al., 2010). Accordingly, we anticipated that CM patients could perform worse in the action prediction task than in the shape prediction task. Furthermore, in this experiment we directly controlled for the influence of intelligent quotient (IQ) on the performance in the two tasks.

**Table 3.** Demographic and clinical information of the three groups of experiment 2.

|  | CM | CND | TD |
| --- | --- | --- | --- |
| *Demographic information* |  |  |  |
| **N (male: female)** | 12:6 | 13:5 | 12:6 |
| **Age in years: mean (SD)** | 12.6 (2.8) | 12.8 (3.4) | 11.1 (2.9) |
| *Clinical information* |  |  |  |
| **IQ: mean (SD)** | 71 (27) | 78 (16) | 107 (10) |
| **Primary malformation** | Molar tooth sign (6); vermis hypoplasia (7); vermis and hemispheres hypoplasia (3); hemispheres hypoplasia (left) and dysplasia (right) (1); rhomboencephalosynapsis (1) | Corpus callosum agenesis (2); thick corpus callosum (1); thalamic hamartoma and UBOs (1); aberrant supracallosal longitudinal bundle (2); none (12) |  |
| **Syndromic/genetic diagnosis** | Joubert syndrome (6); Dandy-Walker malformation (1); unknown (11) | 16p11.2 deletion syndrome (1); SOX5 deletion (1); neurofibromatosis type 1 (1); SCN8A-related disorder (1); Floating-Harbor syndrome (1); unknown (13) |  |

Legend: Legend: CM=Cerebellar malformation; CND=Congenital neurological diseases; TD=Typical development; SD=Standard deviation; IQ=Intelligent quotient; SCN8A=Sodium
voltage-gated channel alpha subunit 8; SOX5=SRY-Box Transcription Factor 5; UBOs=Unidentified bright objects.

As in Experiment 1, the groups were comparable for age (F_2,51_=1.91, p=0.159) and gender (Chi^2^=0.172, p=0.918). In the familiarization phase, the ANOVA revealed significant effect of group (F_2,51_=3.91, p=0.026, η^2^_p_=0.13), showing that CM patients (90.01±1.80%) had lower performance than the other groups (CND: 93.90±1.47%, TD: 94.93±0.98%; all p<0.042). Moreover, a significant effect of task emerged (F_1,51_=19.87, p<0.001, η^2^_p_=0.28), indicating that participants were better at performing the action (95.07±0.98%) than the shape prediction task (90.82±1.85%). However, the group x task interaction was non-significant (F_2,51_=1.30, p=0.282). The follow-up ANCOVA showed a significant effect of the covariate IQ (F_1, 50_=21.70, p<0.001, η^2^_p_=0.3), with better performance in higher IQ individuals (r=0.58), while all other effects were non-significant (all F<3.90, all p>0.053). Thus, performance in the familiarization phase was entirely explained by IQ levels.

The ANOVA on the testing phase revealed significant effectsg of group (F_2,51_=3.44, p=0.040, η^2^_p_=0.12) and probability (F_3,153_=7.38, p<0.001, η^2^_p_=0.13), better qualified by their interaction (F_6,153_=3.36, p=0.004, η^2^_p_=0.12). In detail, TD participants were less accurate in the 10% condition (61.89±6.53%) compared to the 90% (84.75±2.72%; p<0.001) and 60% conditions (80.28±4.14%; p<0.001). In these latter conditions, they also showed higher accuracy compared to the 40% conditions (70.06±6.53%, all p<0.030). In the CND group, a probabilistic facilitation was reliable for the 90% (80.39±4.36%) compared to the 10% condition (70.69±5.23%; p=0.039). Conversely, CM patients’ accuracy was comparable across probability conditions (all p>0.351). Between-groups comparisons showed that, at the highest-probability condition, the two control groups had comparable performance (p=0.455), and both had higher accuracy than CM patients (65.50±4.62%; all p<0.027). For the 60% condition TD participants outperformed CM patients (65.41±3.68%; p=0.027), while no difference emerged for the CND group (75.41±3.68%; all p>0.136). Similarly, for the 40% condition CM patients (61.72±4.25%) were less accurate than CND participants (76.28±4.25%; p=0.033), although the TD group performance did not differ from the other groups (all p>0.218). Conversely, at the lowest-probability condition no difference emerged between groups (CM: 66.22±4.58%). These findings support a domain-general prediction deficit due to cerebellar alterations as expected by the UCT hypothesis (Schmahmann, 2019). The ANOVA also yielded significant effects of task (F_1,51_=4.24, p=0.045, η^2^_p_=0.08), with better performance at the action than shape prediction task, but its interactions with group and probability were non-significant (all p>0.194). Partially in contrast to our hypothesis and with previous research (Cattaneo et al., 2012; Sokolov et al., 2010), this result ruled out that CM patients had worse performance with social stimuli than with physical events. Moreover, in the follow-up ANCOVA, nor the main effect of task neither its interactions were significant (all F<1.65, all p>0.204). Conversely, both the main effect of group (F_2, 51_=4.24, p=0.020, η p=0.15) and the group x probability interaction (F_1, 50_= 18.73, p<0.001, η^2^_p_=0.27) were still significant after partialling out the effects of IQ. Nevertheless, a significant IQ effect was confirmed in this analysis, with higher cognitive levels associated with better performance across conditions (r=0.5). However, interaction effects of IQ with other variables were non-significant (all F<0.75, all p>0.386), suggesting that IQ did not influence the different use of contextual predictions in the three groups. In sum, results showed that neither in the action prediction task nor in the shape prediction task did CM patients present a contextual modulation effect, while the control groups showed a reliable use of contextual priors in both tasks (Fig. 4).

**Figure 4.**
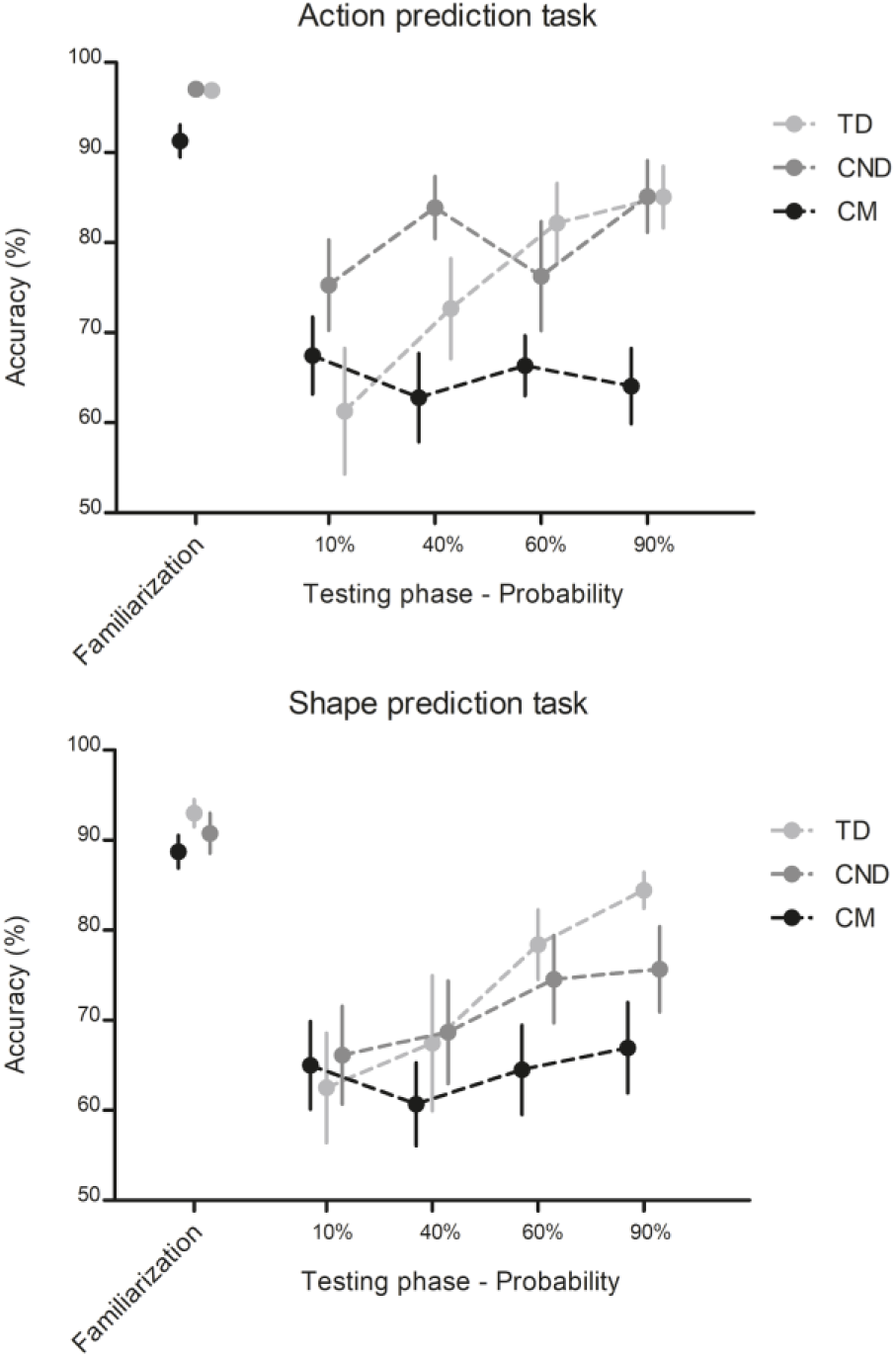
Results of Experiment 2. Eighteen patients per group were administered with the action prediction task and the shape prediction task. Data processing and statistical analyses were performed following the same design of Experiment 1 with the addition of task as within-subject variable. Then, two follow-up ANCOVAs were used to partial out the effects of IQ. In the familiarization phase, the ANCOVA showed a significant effect of the covariate IQ (p<0.001), while all other effects were nonsignificant (all p>0.053). In the testing phase, the main effect of group (p=0.020) and the group x probability interaction (p<0.001) were still significant after partialling out the effects of IQ, but nor the main effect of task neither its interactions were significant (all p>0.204).

### Experiment 3: Effects of the VR social prediction training on the use of contextual prediction

In this experiment, we tested how social prediction can be boosted in CM patients by a VR training specifically developed to improve predictive abilities in a social scenario (VR-SPIRIT; (Butti, Biffi, et al., 2020)). In the VR-SPIRIT, the participants were immersed in a playground scenario and, in each of the 80 total trials, they were asked to compete with one of four avatars for reaching one of three recreational objects. Specific features of the scenario forced the participants to anticipate the behavioral preference of each avatar, which, crucially, was associated to the objects with pre-established probabilities. The use of predictive and random strategies in anticipating avatars’ intentions was computed respectively, by the mean percentage of scores obtained when the probabilistic avatar-object association gave clues on avatar’s intention (prediction score) and by the mean percentage of scores obtained when context did not provide reliable information (random score). CM patients were randomly assigned to the VR-SPIRIT or to a VR-based motor training and then exposed for eight 45-minute sessions to one of these two interventions, which were run on the same VR platform. Before and after the training, CM patients were administered a VR evaluation session exploiting the same probabilistic design of the training sessions, but presenting a different scenario, and the action prediction task adopted in the former experiments. This way, we evaluated the transferability of a VR experience to the use of contextual priors for predicting social events in patients with cerebellar alterations, paving the way to new rehabilitative approaches for CCAS (Argyropoulos et al., 2020).

Preliminary analyses confirmed that participants assigned to the VR-SPIRIT and to the motor training were comparable for age (Z=-1.29, p=0.199), cognitive level (Z=-0.341, p=0.733), and gender (Chi^2^=0.267, p=0.606). For the pre-training assessment, the analyses did not reveal any significant difference between groups, either in the action prediction task (all Z<|1.22|, all p>0.246) or in the VR evaluation session (all Z<|0.68|, all p>0.495). Considering the within-group changes after the intervention, only the experimental group improved performance in the VR evaluation scenario, particularly in using predictive strategies when the context gave clues on avatars’ intentions (Z=-2.81, p=0.005, r=0.89), but not in the random score (Z=-0.120, p=0.905). These results could not be accounted for by a diverse exposition to the VR environment, since also motor training-participants increased their abilities in moving on the VR platform as revealed by shortened trial duration (Z= – 2.60, p=0.009, r=0.82). However, this latter group did not present significant changes in other variables of the VR sessions (all Z<| 1.86|, all p>0.062). For the action prediction task, Wilcoxon tests did not yield any significant change (all Z<|1.17|, all p>0.241). Crucially, after the training, the VR-SPIRIT group performed significantly better than the motor-training group in the VR evaluation session, with higher total score (Z=-3.12, p=0.002, r=0.70) and prediction score (Z=-2.81, p=0.025, r=0.63). Furthermore, the beta index at the action prediction task was higher after the VR-SPIRIT than after the motor training (Z= −1.97, p= 0.049, r= 0.44), thus suggesting that training predictive abilities in a social VR scenario also boosted the implicit learning and use of contextual priors during action perception (Fig. 5 and Table 4).

**Figure 5.**
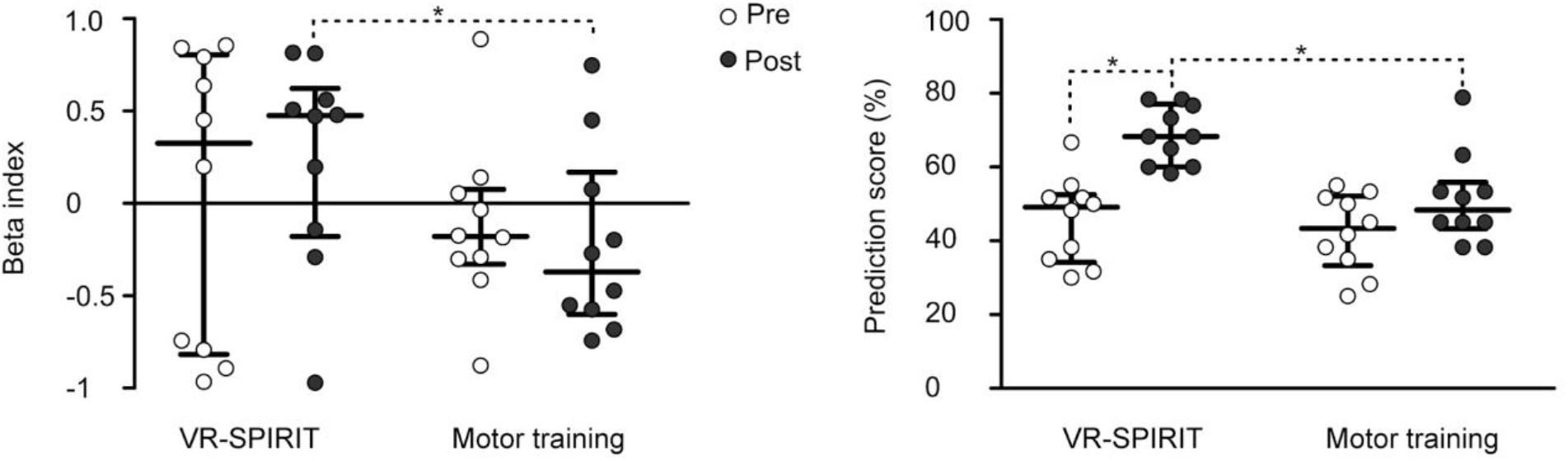
Results of Experiment 3. Beta index of the action prediction task and percentage of prediction scores in the VR evaluation session for the two groups (N = 10 per group) before and after the rehabilitative training. Mann-Whitney U tests revealed that, after the trainings, the VR-SPIRIT group compared to the motor training group was more able to use contextual information to predict other’s behavior in both the action prediction task (p=0.049) and the VR session (p=0.025). Wilcoxon signed-ranks indicated that only the VR-SPIRIT group showed significant improvements in the prediction scores of the VR session (p=0.005). Long bars indicate median, short bars indicate interquartile range; circles indicate individual performance; asterisks indicate significant comparisons.

**Table 4.** Between-groups comparisons for the action prediction task and the VR evaluation sessions before and after the two rehabilitative interventions. Significant results are reported in bold.

|  |  | **Experimental training** | **Control training** | **Mann-Whitney U** |
| --- | --- | --- | --- | --- |
|  |  | Median (IQR)/Z (p) | Median (IQR)/Z (p) | Z (p) |
| *Action prediction task* |  |  |  |  |
| **Familiarization (%)** | Pre | 96 (8) | 90 (3) | -1.22 (0.223) |
|  | Post | 93 (8) | 91 (10) | -0.91 (0.363) |
| **Beta index** | Pre | 0.33 (1.53) | -0.18 (0.33) | -0.45 (0.650) |
|  | Post | 0.48 (0.60) | -0.37 (0.57) | **-1.97 (0.049)** |
| *VR Evaluation session* |  |  |  |  |
| **Total score** | Pre | 36 (12) | 31 (9) | -0.54 (0.591) |
|  | Post | 49 (6) | 36 (9) | **-3.12 (0.002)** |
| **Trial duration (sec)** | Pre | 6.5 (1.1) | 6.4 (0.4) | -0.23 (0.821) |
|  | Post | 5.8 (0.3) | 6.0 (0.4) | -1.74 (0.082) |
| **Prediction score (%)** | Pre | 49 (16) | 43 (15) | -0.53 (0.596) |
|  | Post | 68 (15) | 48 (8) | **-2.81 (0.025)** |
| **Random score (%)** | Pre | 28 (10) | 28 (13) | -0.50 (0.617) |
|  | Post | 30 (11) | 38 (14) | -1.04 (0.298) |

Legend: IQR=Interquartile range; VR=Virtual reality

## Discussion

In this work, we tested the effects of cerebellar alterations on the predictive processing of incoming social and physical events. In line with the UCT hypothesis (Schmahmann, 1996, 2019), our results showed that cerebellar patients showed a domain-general prediction deficit. In a recent ALE meta-analysis, Siman-Tov and colleagues found a consistent activation of the cerebellum during predictive tasks assessing multiple domains (Siman-Tov et al., 2019). The authors proposed that, through its role in sequencing (M. Leggio & Molinari, 2015) and error-based learning (Peterburs & Desmond, 2016), the cerebellum may contribute to a wide cortical-subcortical network underlying domain-general predictions. Here, we further elucidated this contribution by showing that cerebellar alterations impaired the processing of contextual information, resulting in less precise predictions when the available sensorial information was not sufficient to inform on the incoming events (Friston, 2012a). The anatomo-functional organization of the cerebellum seems optimal for processing the cooccurrence between contextual and sensorial cues and integrating them into internal prediction models (Ishikawa et al., 2016). After the exposition to repetitive context-event associations, this predictive processing would result in contextual priors that are matched with bottom-up sensorial inputs, crucially contributing to selecting the most probable event within a specific context. Thus, within a Bayesian paradigm of brain functioning, the cerebellum could modulate the interactions between cortical nodes of specific cognitive networks by supplying contextual priors that constrain stimulus processing at any stage (Sokolov et al., 2017).

Our result of comparable deficits in the prediction of social and physical events suggests that the cerebellum may apply its predictive computation regardless of the social nature of the processed information. This finding is partially in contrast to a previous study on adult patients with acquired cerebellar damage (M. G. Leggio et al., 2008), in which a general cognitive sequencing impairment was reported for both actions and abstract figures. However, the authors found associations between lesions to the left and right hemispheres and, respectively, the processing of pictorial and verbal stimuli, in line with the hypothesis of a universal computation exerted on diverse information by specific cortico-cerebellar loops (Stoodley & Schmahmann, 2010). Thus, we could speculate that the lack of task differences might reflect the complex nature of cerebellar malformations presented by our sample (Poretti et al., 2016), which did not allow disentangling specific anatomo-functional associations. Furthermore, congenital alterations could affect the development and functioning of the cerebellum differently from acquired damage, also due to developmental diaschisis effects on other cortical areas (Stoodley & Limperopoulos, 2016).

Despite the cerebellar coding of context-based predictions may operate across different domains, an impairment in using contextual priors is likely to have major consequences in the social domain, in which the context is mostly crucial to disambiguate others’ behavior (Brown & Brüne, 2012). Accordingly, we showed that CM patients were more impaired than CND patients not only in using contextual priors to predict the unfolding of others’ actions, but also in discriminating the most likely facial expression within a specific social scenario, as assessed by the NEPSY-II non-verbal ToM subtest. Furthermore, the context-based prediction abilities of CND patients were associated with their performance at the NEPSY-II subtests assessing affect recognition and non-verbal ToM abilities. Thus, impairments in using contextual priors due to cerebellar defect disrupted low-level inferences of motor-intentions (Baker et al., 2005) as well as higher-level predictions involved in understanding others’ emotions and mental states (Koster-Hale & Saxe, 2013).

Through its connections with cortical areas engaged in action perception (Sokolov et al., 2010), affect processing (Adamaszek et al., 2017) and mentalization (Van Overwalle & Mariën, 2016), the cerebellum could provide priors embedding contextual information, modulating these cortical networks and crucially contributing to select the best matching between external information and internal expectations (Van Overwalle et al., 2019). Accordingly, a recent study reported that diminished functional connectivity between the cerebellum and brain areas activated in social tasks was associated with deficits of social cognitive processing at different levels (Clausi et al., 2019). Thus, our findings supported the hypothesis that cerebellar alterations could result in a weaker or absent reliance on contextual information to understand others’ intentions (Butti, Corti, et al., 2020), ultimately leading to the social perception deficits documented in cerebellar patients (Hoche et al., 2016; Schmahmann et al., 2007).

Although the clinical relevance of social perception deficits has long been described (Schmahmann & Sherman, 1998; Tavano et al., 2007; Van Overwalle et al., 2020) and further confirmed by our findings, only few studies have so far proposed treatments for CCAS symptoms (Gagliardi et al., 2015; T. Ito et al., 2010; Maeshima & Osawa, 2007; Ruffieux et al., 2017) and none of them have targeted social cognition. Here, we reported first evidence that a VR training specifically designed to target social prediction abilities in CM patients may improve the use of predictive strategies in understanding others’ behavior and enhance the reliance on contextual information to predict ongoing actions. A recent task-force paper has proposed that rehabilitation should aim to make cerebellar patients aware of their deficits so that implicit, automatic cerebellar functions could be compensated by explicit thought processes, acting as an “external cerebellum” (Argyropoulos et al., 2020; Schmahmann, 2010). Accordingly, we could argue that VR-SPIRIT trained participants in learning explicit context-behavior associations and that this improvement was at least partially generalizable to the use of implicit, context-based predictions during the action prediction task. These results encourage the testing of how the improvements of context-based action predictions after the VR-SPIRIT may transfer to more general social cognition abilities. Moreover, combining the VR-SPIRIT with non-invasive stimulation of the cerebellum (Oldrati & Schutter, 2018) to promote the neural plasticity of cerebellar circuits underpinning social cognition (Boggio et al., 2015) might further boost the generalization of the social prediction improvements.

The results of this study should be discussed considering its limitations. Even though we calculated a-priori the sample size for Experiments 1 and 2, we cannot exclude that the absence of differences in executing the two prediction tasks might reflect a power issue also due to the heterogeneity of our clinical groups. The complex nature of the congenital diseases presented by both CM and CND samples prevented us also from verifying the impact of the localization and extension of the cerebellar malformation on the prediction of social and non-social stimuli. Moreover, our experiments did not allow us to clarify whether cerebellar alterations affected the building or the use of contextual priors. Despite a deficit in encoding priors is in line with the cerebellar role in sequencing (M. G. Leggio et al., 2008; M. Leggio & Molinari, 2015) and implicit learning of contextual regularities (Bellebaum & Daum, 2011; Ulasoglu-Yildiz & Gurvit, 2019), we could not exclude that CM patients may have encoded the context-event associations, but then they could not use these contextual priors in condition of perceptual uncertainty (Sokolov et al., 2017). Regarding the effects of the VR-SPIRIT in Experiment 3, the small number of recruited patients and the limited choice of outcome measures asks for caution in generalizing the results. However, the results point to the validity of this training approach for the social cognition domain by highlighting the transferability of its effects on the use of contextual priors for action prediction.

In conclusion, the present study provided evidence that a general deficit in using contextual priors to predict incoming sensorial information might underlie the social perception deficits reported in patients with cerebellar alterations. However, not only did we document these deficits and probe their relevance for general social cognition abilities; we also provided evidence that a short-lasting, but intense (i.e., eight sessions in two weeks), VR training designed to boost the learning of statistical regularities in others’ behavior can reinforce context-based predictions across different scenarios. This paves the way for new approaches to the rehabilitation of patients with CCAS specifically based on the predictive computational mode of the cerebellum.

## Materials and Methods

### Participants

Pediatric patients were recruited at the Unit of neuropsychiatry and neurorehabilitation of the Scientific Institute Medea. For the group with cerebellar malformations (CM), inclusion criteria were: i) diagnosis of a non-progressive congenital cerebellar malformation; ii) age ranging from 7 to 18 years; iii) absence of severe cognitive delay, with a Full-Scale Intelligent Quotient (FSIQ) ≥ 40 at the age-corresponding Wechsler scale; iv) absence of severe motor and visual impairments that could interfere with task execution. A full description of the enrolled sample in the study is reported in Table 5.

**Table 5.**
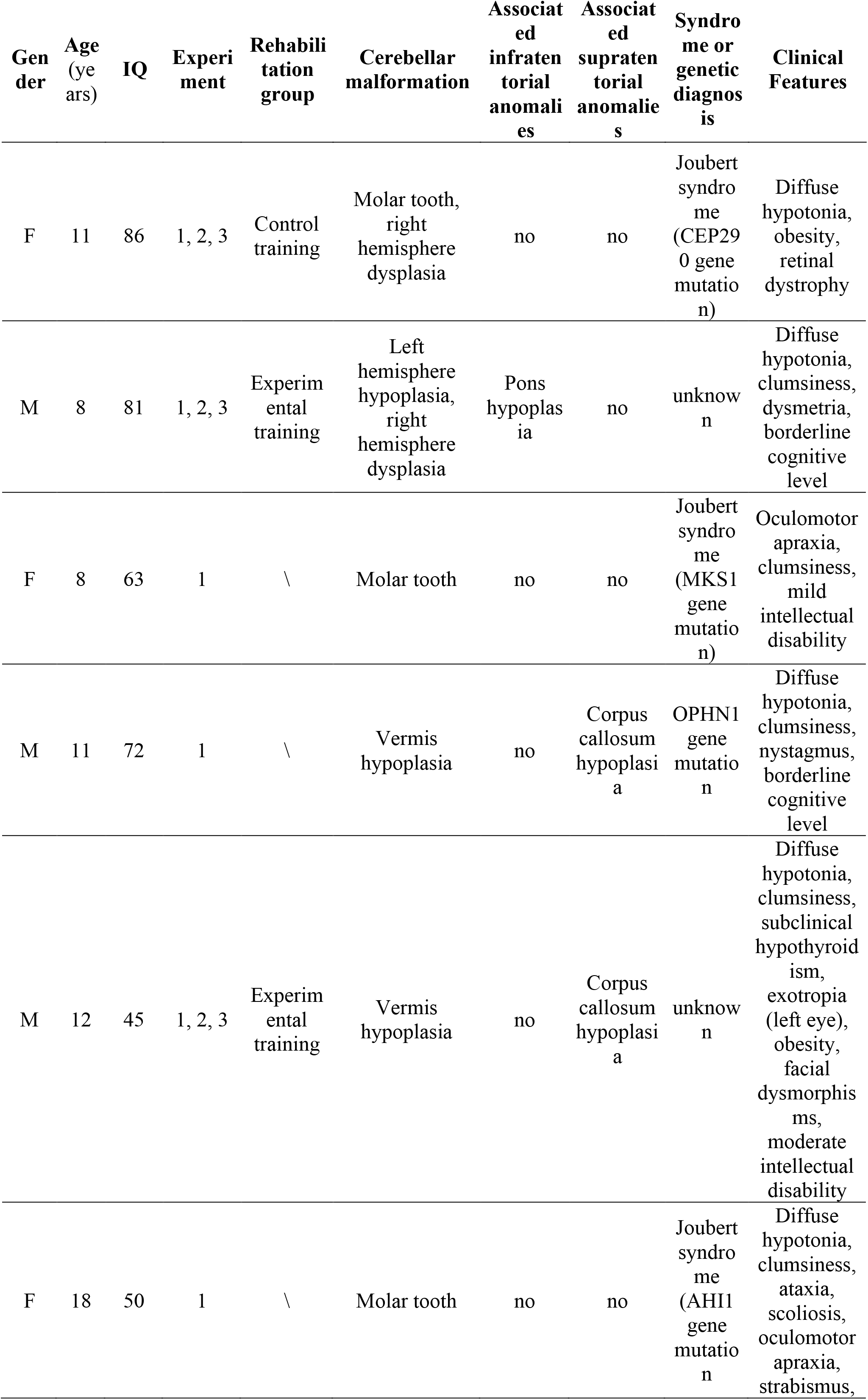
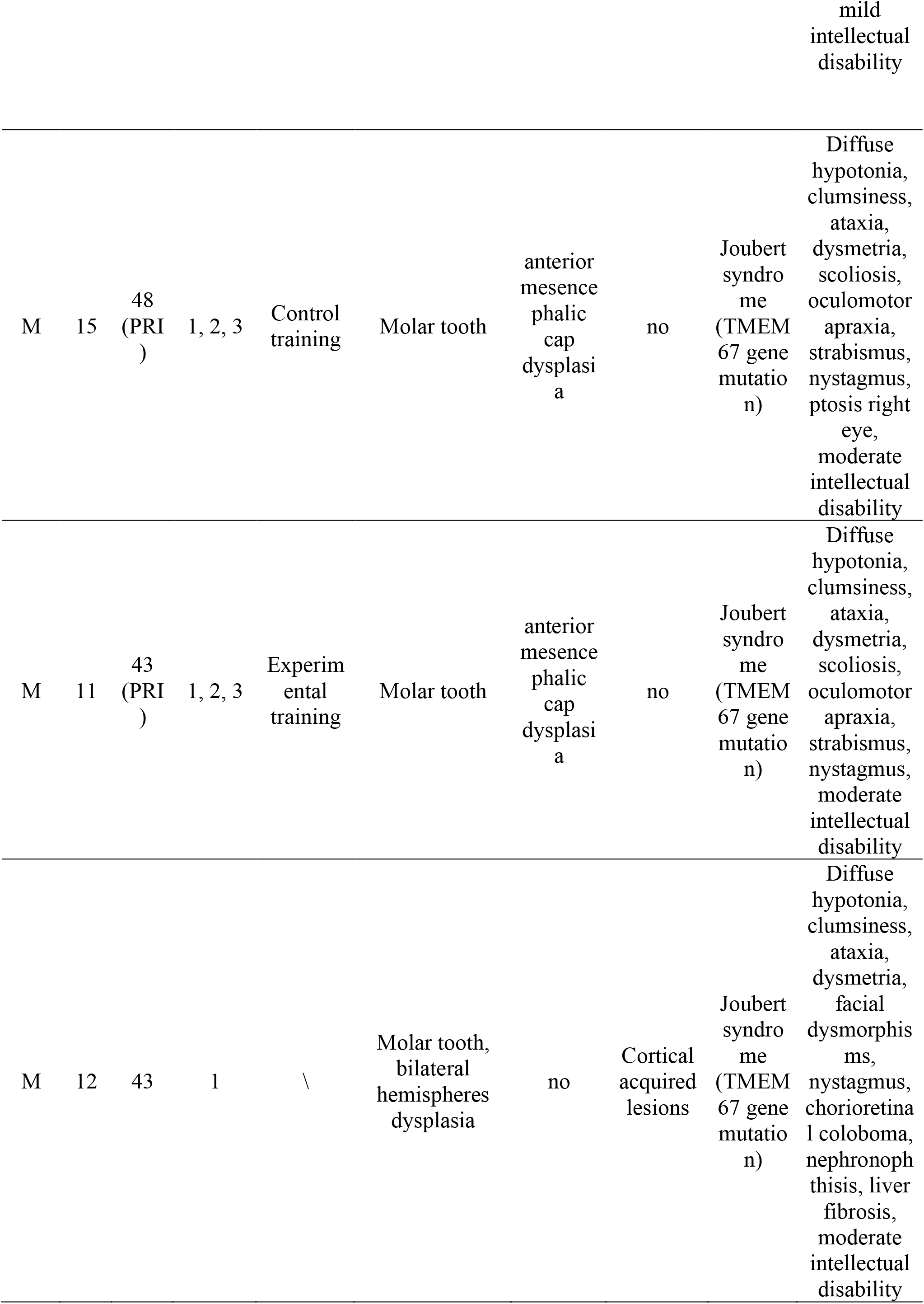
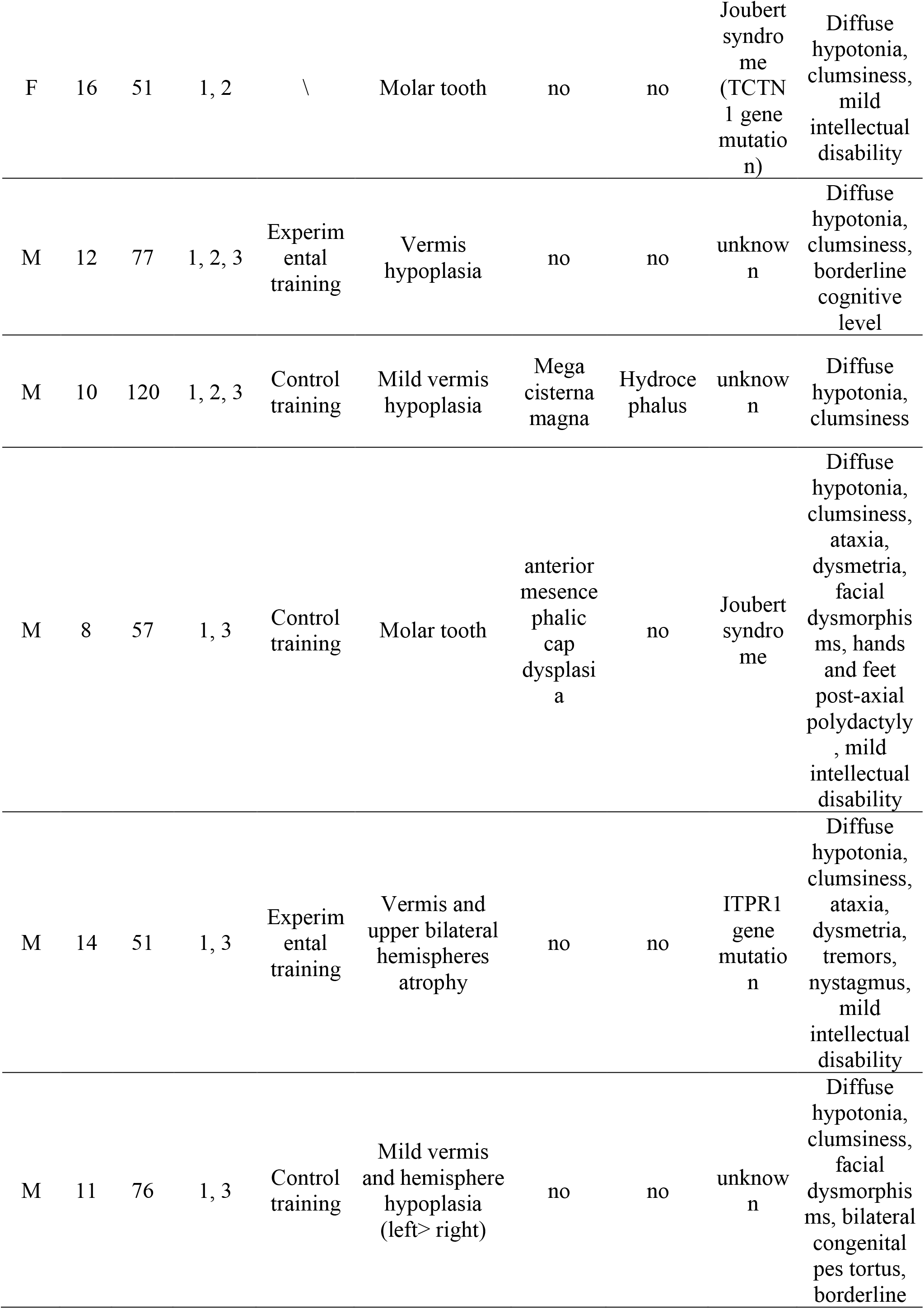
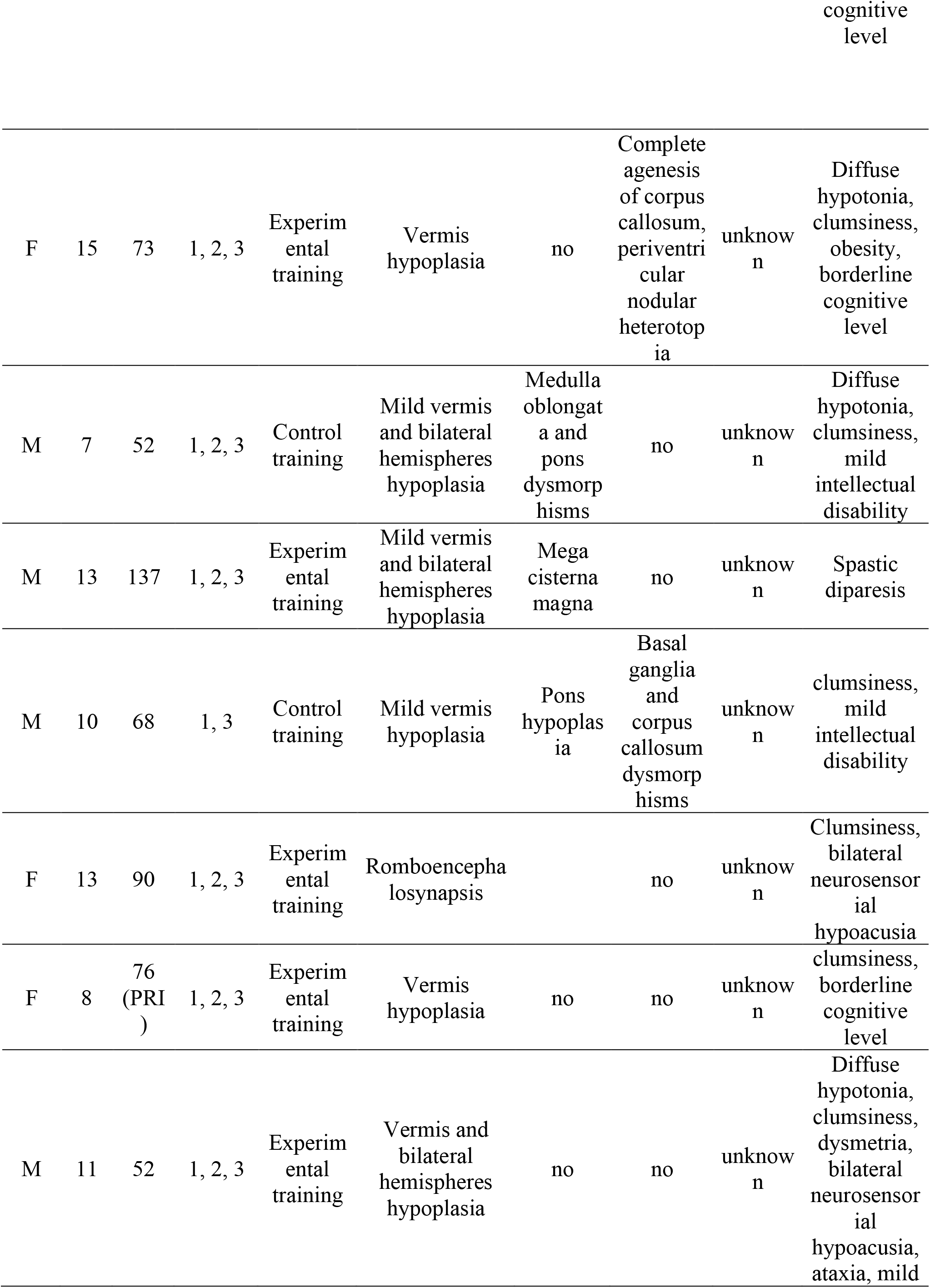
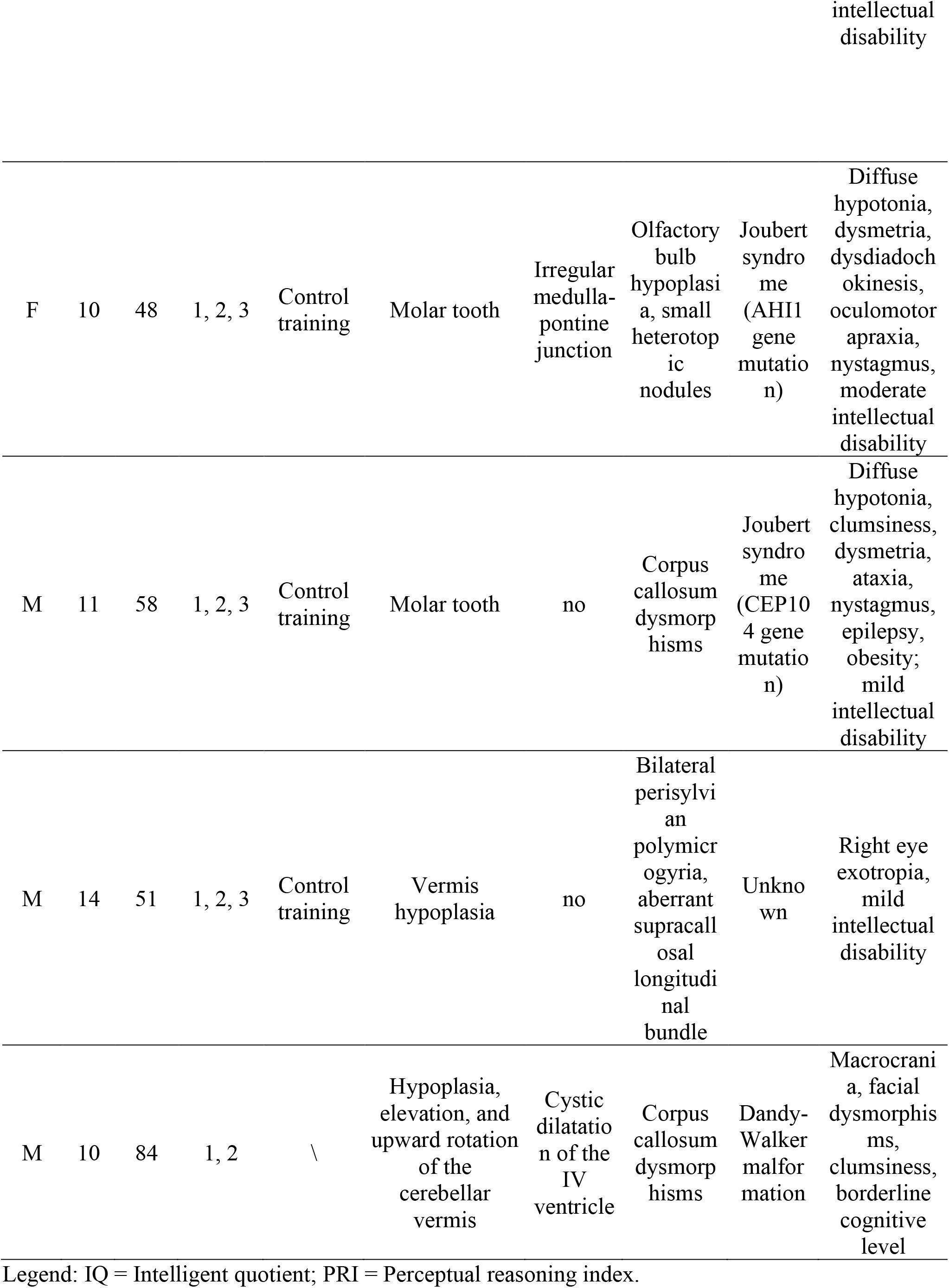
Neuroradiological and clinical information of the group with cerebellar malformation.

For the control group of patients with congenital neurological disorders (CND), we adopted the same inclusion criteria but enrolled patients without cerebellar malformation as revealed by MRI reports (Table 6).

**Table 6.**
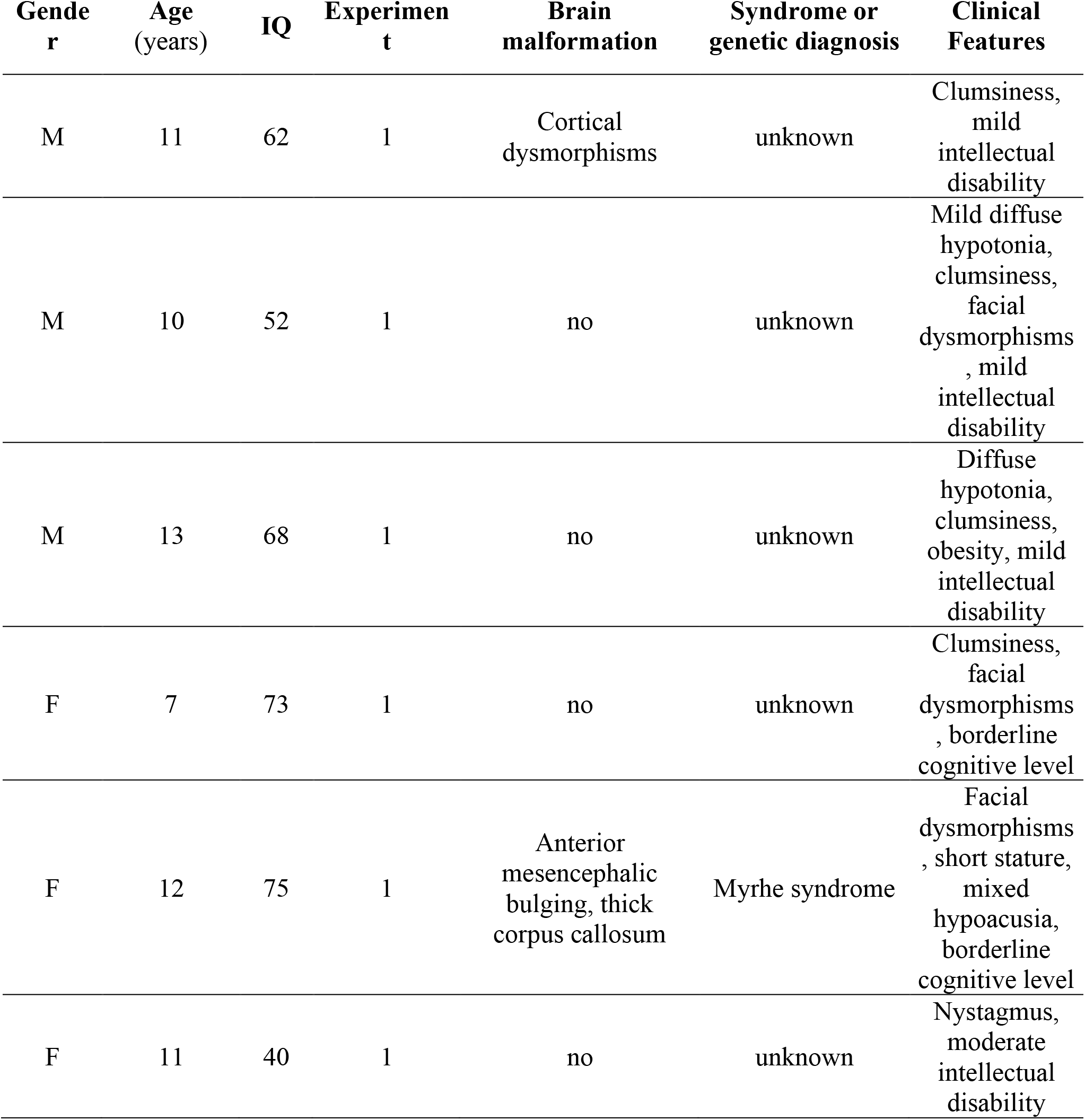
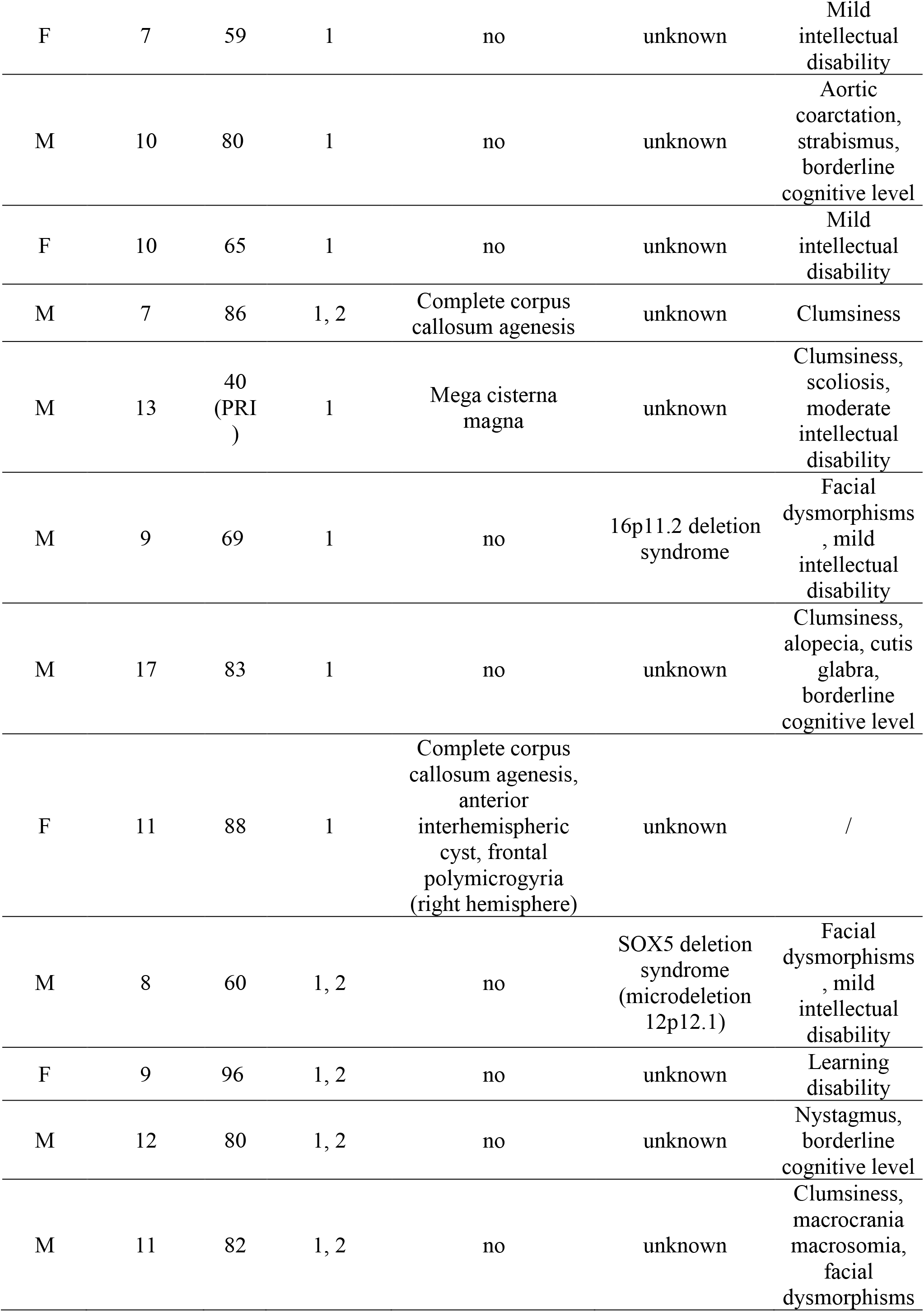
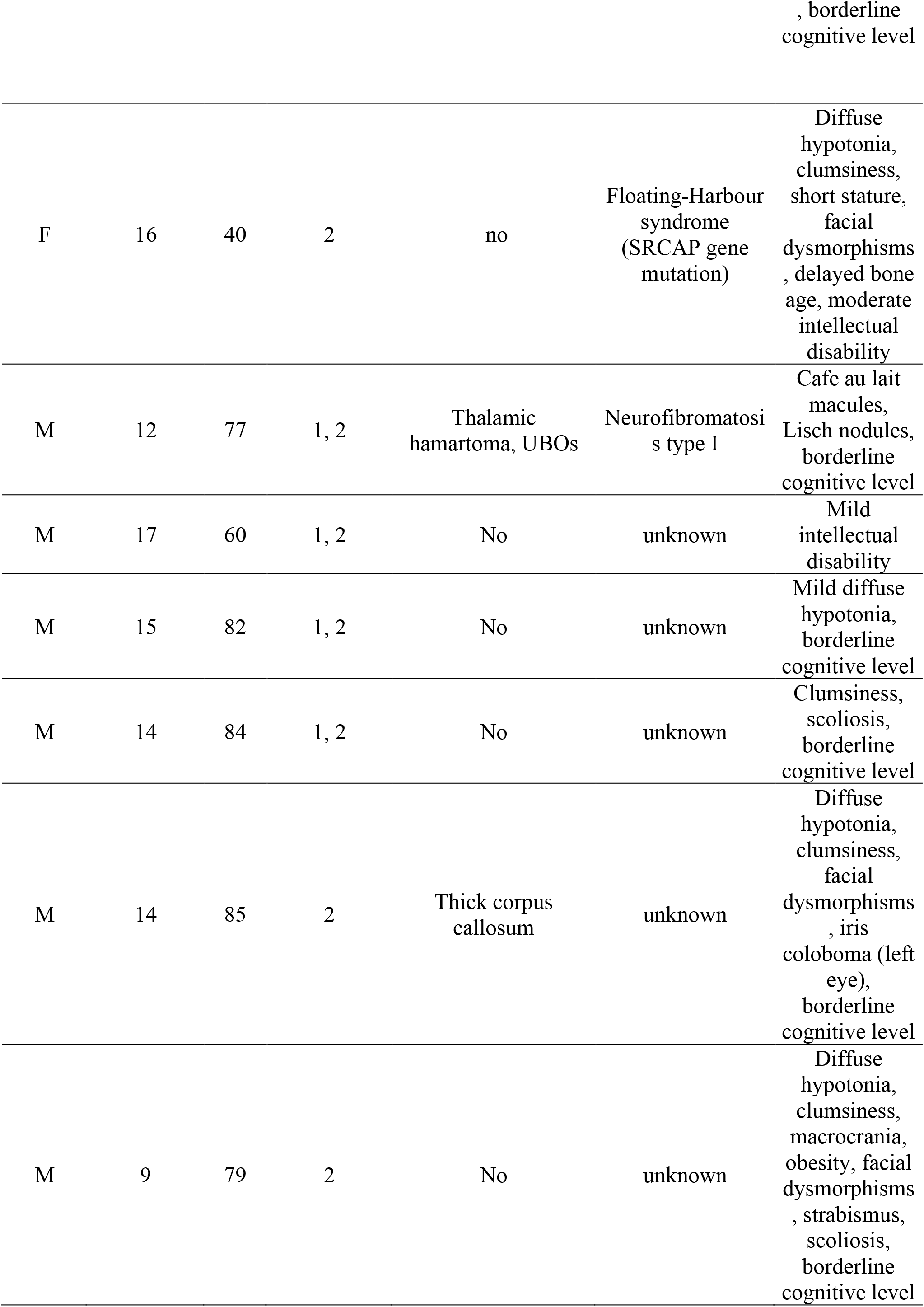
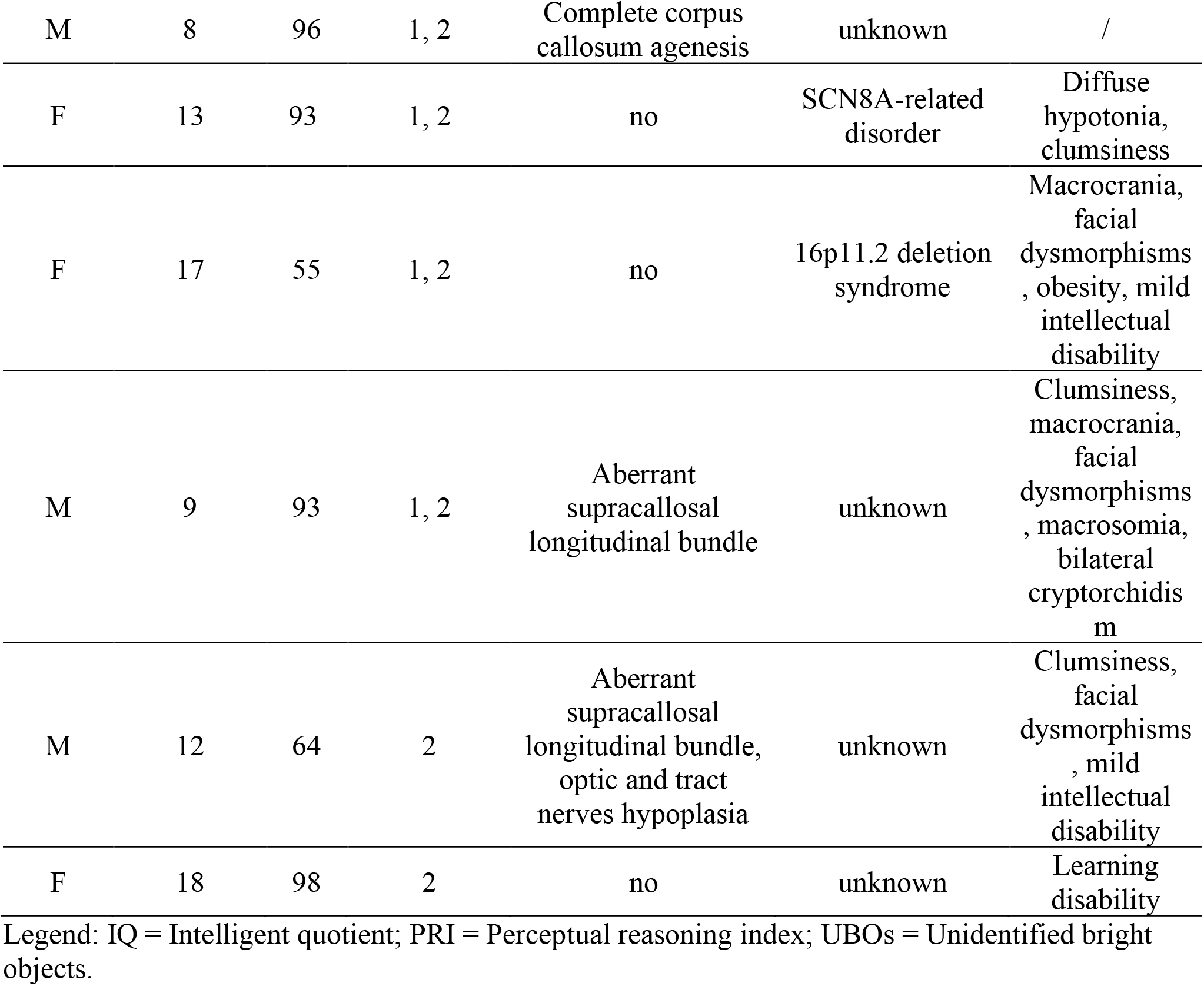
Neuroradiological and clinical information of patients with congenital neurological disorders not affecting the cerebellum.

| **Gende**<br>**r** | **Age**<br>**(years)** | **IQ** | **Experimen**<br>**t** | **Brain**<br>**malformation** | **Syndrome or**<br>**genetic diagnosis** | **Clinical**<br>**Features** |
| --- | --- | --- | --- | --- | --- | --- |
| M | 11 | 62 | 1 | Cortical<br>dysmorphisms | unknown | Clumsiness,<br>mild<br>intellectual<br>disability |
| M | 10 | 52 | 1 | no | unknown | Mild diffuse<br>hypotonia,<br>clumsiness,<br>facial<br>dysmorphisms<br>, mild<br>intellectual<br>disability |
| M | 13 | 68 | 1 | no | unknown | Diffuse<br>hypotonia,<br>clumsiness,<br>obesity, mild<br>intellectual<br>disability |
| F | 7 | 73 | 1 | no | unknown | Clumsiness,<br>facial<br>dysmorphisms<br>, borderline<br>cognitive level |
| F | 12 | 75 | 1 | Anterior<br>mesencephalic<br>bulging, thick<br>corpus callosum | Myrhe syndrome | Facial<br>dysmorphisms<br>, short stature,<br>mixed<br>hypoacusia,<br>borderline<br>cognitive level |
| F | 11 | 40 | 1 | no | unknown | Nystagmus,<br>moderate<br>intellectual<br>disability |
| F | 7 | 59 | 1 | no | unknown | Mild intellectual disability |
| M | 10 | 80 | 1 | no | unknown | Aortic coarctation, strabismus, borderline cognitive level |
| F | 10 | 65 | 1 | no | unknown | Mild intellectual disability |
| M | 7 | 86 | 1, 2 | Complete corpus callosum agenesis | unknown | Clumsiness |
| M | 13 | 40 (PRI) | 1 | Mega cisterna magna | unknown | Clumsiness, scoliosis, moderate intellectual disability |
| M | 9 | 69 | 1 | no | 16p11.2 deletion syndrome | Facial dysmorphisms, mild intellectual disability |
| M | 17 | 83 | 1 | no | unknown | Clumsiness, alopecia, cutis glabra, borderline cognitive level |
| F | 11 | 88 | 1 | Complete corpus callosum agenesis, anterior interhemispheric cyst, frontal polymicrogyria (right hemisphere) | unknown | / |
| M | 8 | 60 | 1, 2 | no | SOX5 deletion syndrome (microdeletion 12p12.1) | Facial dysmorphisms, mild intellectual disability |
| F | 9 | 96 | 1, 2 | no | unknown | Learning disability |
| M | 12 | 80 | 1, 2 | no | unknown | Nystagmus, borderline cognitive level |
| M | 11 | 82 | 1, 2 | no | unknown | Clumsiness, macrocrania macrosomia, facial dysmorphisms |
| F | 16 | 40 | 2 | no | Floating-Harbour syndrome (SRCAP gene mutation) | Diffuse hypotonia, clumsiness, short stature, facial dysmorphisms, delayed bone age, moderate intellectual disability |
| M | 12 | 77 | 1, 2 | Thalamic hamartoma, UBOs | Neurofibromatosis type I | Cafe au lait macules, Lisch nodules, borderline cognitive level |
| M | 17 | 60 | 1, 2 | No | unknown | Mild intellectual disability |
| M | 15 | 82 | 1, 2 | No | unknown | Mild diffuse hypotonia, borderline cognitive level |
| M | 14 | 84 | 1, 2 | No | unknown | Clumsiness, scoliosis, borderline cognitive level |
| M | 14 | 85 | 2 | Thick corpus callosum | unknown | Diffuse hypotonia, clumsiness, facial dysmorphisms, iris coloboma (left eye), borderline cognitive level |
| M | 9 | 79 | 2 | No | unknown | Diffuse hypotonia, clumsiness, macrocrania, obesity, facial dysmorphisms, strabismus, scoliosis, borderline cognitive level |
| M | 8 | 96 | 1, 2 | Complete corpus callosum agenesis | unknown | / |
| F | 13 | 93 | 1, 2 | no | SCN8A-related disorder | Diffuse hypotonia, clumsiness |
| F | 17 | 55 | 1, 2 | no | 16p11.2 deletion syndrome | Macrocrania, facial dysmorphisms, obesity, mild intellectual disability |
| M | 9 | 93 | 1, 2 | Aberrant supracallosal longitudinal bundle | unknown | Clumsiness, macrocrania, facial dysmorphisms, macrosomia, bilateral cryptorchidism |
| M | 12 | 64 | 2 | Aberrant supracallosal longitudinal bundle, optic and tract nerves hypoplasia | unknown | Clumsiness, facial dysmorphisms, mild intellectual disability |
| F | 18 | 98 | 2 | no | unknown | Learning disability |

Legend: IQ = Intelligent quotient; PRI = Perceptual reasoning index; UBOs = Unidentified bright objects.

For Experiment 1, a total of 26 children and adolescents with CM and an equal number of peers with CND were recruited. Furthermore, 26 healthy peers with typical development (TD) and no previous history of any neurological or psychiatric disorder were recruited at local schools. For Experiment 2, a total of 18 participants per group were enrolled. Raven’s progressive matrices (Raven, 1982) were administered to TD participants to obtain a measure of their cognitive level comparable with the FSIQ of the two clinical groups (Mungkhetklang et al., 2016). For Experiment 3, 20 CM patients were assigned to one of the two training conditions according to a stratified permuted block randomized procedure (see Butti, Biffi et al., 2020 for further details).

All participants were asked to provide their informed consent to the study and their parents signed a written informed consent before starting each experiment. All procedures were conducted in accordance with the Declaration of Helsinki and approved by the local ethical committee (Prot. N.34/18 – CE for Experiments 1,2; Prot. N.17/18 – CE for Experiment 3).

### Action prediction task (Experiments 1, 2, 3)

We adopted the same task of Amoruso and colleagues (Amoruso *et al.*, 2019). In this two-alternative forced choice (2AFC) task, participants were exposed to short videos that depicted a child executing distinct grasping actions with two diverse intentions and they were asked to predict the final outcome of the action (i.e., the motor intention). Before starting the experiment, participants were introduced to the objects displayed in the videos, namely an apple and a glass, and were informed about the different feasible object-manipulations associated with either individual (i.e., to eat/drink) or interpersonal actions (i.e., to offer). Importantly, the apple and the glass were presented, respectively, on a plate and on a tablecloth that could be of two different colors (i.e., orange or violet for the plate, white or blue for the tablecloth).

The task consisted of two blocks, each comprising a familiarization phase (80 trials) immediately followed by a testing phase (40 trials), for a total of 240 trials. Participants were presented with the same videos in both phases, but in the testing one video duration was drastically reduced. In the familiarization phase, videos were stopped just two frames before the hand-contact with the object (25 frames, each frame lasting 33.33 ms for a total of 833 ms), thus displaying almost in full the action unfolding. Importantly, during this phase the probability of presentation of a given action in association with a specific contextual cue was biased by setting the probability of action-contextual cue associations to 10%, 40%, 60%, or 90% of the total number of trials. In this way, the prior expectation was implicitly manipulated (for further details see (Amoruso *et al.,* 2019)). The type of action-context association was counterbalanced between participants but remained constant in the two blocks of familiarization within the same participant.

In the testing phase, video duration was drastically shortened (500 ms, 15 frames), thus hindering kinematic information. Given the manipulation of the probability of co-occurrence between the actions’ intentions and the color of contextual cues in the familiarization phase, we expected that participants could implicitly rely on the contextual priors to overcome kinematics uncertainty, presenting a probabilistic modulation in their responses. Notably, differently from the familiarization phase, during the testing phase each of the 8 videos was presented for an equal number of trials (i.e., five trials in each blocks). Moreover, no information about the probability of associations was explicitly given to participants.

For both phases, each trial started with the presentation of a fixation cross lasting 2,000 ms, followed by video presentation. Immediately after or during the videos (for the familiarization or the testing phase, respectively) the Italian descriptors of the two possible actions (i.e., the verbs *“mangiare”* or “*bere*” and *“offrire”,* in English “to eat” or “to drink” and “to offer”) were presented until a response was recorded. The descriptors, written in white on a black background, were located on the right and on the left side of the screen, with the location counterbalanced between participants and consistent across trials and blocks for each participant. Participants had to respond by pressing with their right or left index finger, respectively, the “m” (right) or the “z” (left) computer key of a QWERTY keyboard. The experiment was ran in a single session lasting ~40 min, using E-Prime V2 software (Psychology Software Tools, Inc., Pittsburgh, PA, United States) on a 15.4-inch LCD computer screen (resolution 1600 × 900 pixels, refresh rate 60 Hz). Participants were asked to sit in front of the monitor at 60 cm. Short breaks were allowed between blocks and phases.

### Shape prediction task (Experiment 2)

This 2AFC task was developed by following the same administration procedure and structure of the action prediction task, but moving geometrical shapes were instead displayed. Participants were asked to observe videos depicting one of four possible two-dimensional geometric shapes moving from the left side of the screen toward a complementary receptor figure placed at the center of the screen. The moving shape could be either right-angle polygons (i.e., a square or a rectangle), or acute angles polygons (i.e., a parallelogram or a trapezoid). Crucially, within each couple of polygons, one was equal length sided (i.e., the square or the parallelogram), while the other was unequal length sided (i.e., the rectangle or the trapezoid). Accordingly, the complementary receptor figure presented, along its left side, a concavity serving as binding site in which the right side of the moving shape could fit. This concavity could be either right-angle or acute-angle shaped according to the type of polygon displayed in the trial. Thus, the right-angle or the acute-angle nature of the polygon was known since the beginning of the trial, while participants had to discriminate between the equal vs. the unequal length sided polygon in each pair. Since the shapes of each pair of polygons looked similar on their right side, the identity of the specific polygon could be detected with the increased visibility of the horizontal segments during the movement, according to the ratio between the major and minor axes of the figure.

Before starting the experiment, participants were introduced to the stimuli displayed in the videos through the presentation of paper-made reproductions of the four polygons and they were informed about the differences in segment length. To simplify the response modality as compared to the original version for adults (Bianco *et al.,* 2020), here we qualified the equal length sided polygons as “short” and the unequal length side polygons as “long”. Thus, for both the familiarization and the testing phases, participants were asked to report whether the moving shape was a short (i.e., square or parallelogram) or a long (i.e., rectangle or trapezoid) shape. Immediately after the videos, the Italian descriptors of the two possible answers (i.e., *“corta”* o “*lunga*”, in English “short” or “long”) were presented in the prompt response frame.

Importantly, each couple of polygons and the respective receptor could be differently colored, using the same probabilistic manipulation as for the action perception task (Figure 2). During the familiarization phase, the shapes fully appeared on the screen so that they could be easily identified (933 ms, corresponding to 28 frames), while, in the testing phase, videos were interrupted one frame after the halfway appearance of the horizontal segment (500 ms, corresponding to 15 frames), thus providing minimal information about the specific shape and prompting the use of the contextual priors to overcome sensory uncertainty.

A resume of the two predictive tasks is reported in Fig. 1.

### VR rehabilitative interventions (Experiment 3)

The VR sessions were administered at the Grail Lab (Motek, Amsterdam, NL) at the Scientific Institute, IRCCS E. Medea. In the experimental VR-SPIRIT, participants were immersed in a playground scenario including a swing, a circular carousel and a rocking carousel. In each trial, participants were asked to compete with one of four avatars for reaching one of these objects. To do so, the participants had to anticipate the behavioral preference of each avatar, since they could not pass the avatar after the direction of its movement was clear. Crucially, the avatars were associated with pre-established probabilities to the objects, so that each of three avatars moved toward his preferred object in the 80% of trials and chose each of the other two objects in the 10% of trials. Conversely, the fourth avatar moved to each object with the same probability. The behavioral preferences of the avatars remained constant within each session (80 trials), so that the participants could learn the probabilistic avatar-object associations, but they were changed across sections.

The active control training consisted in a navigational game and in four GRAIL games previously used in motor rehabilitation (Cesareo *et al.,* 2017). Notably, no social agents were presented, and no prediction abilities were required while playing these games.

Eight 45-minute sessions of one of the two rehabilitative interventions were administered to each participant (for more details see Butti *et al.,* 2020*a*).

### Data handling and statistical analyses

For Experiment 1 we calculated a total sample size of 78 subjects, considering a 4 probabilities x 3 groups mixed-model ANOVA and expecting a medium effect size of f (U)=0.25 as reported by a previous study using the same task on pediatric patients (Amoruso et al., 2019), with an alpha level of 0.05 and a power of 0.80 (1-Beta). Accordingly, we enrolled 26 patients per group. Firstly, we adopted a one-way ANOVA and a Chi-square test to verify that groups were comparable for age and gender. For the action prediction task, response times (RT) were recorded, but not included in the analyses due to the likely impact of motor difficulties within the two clinical groups. However, we excluded trials with anticipated or out-of-time responses (RT<150 ms or >5,000 ms) and then calculated the percentage of correct responses (accuracy) in the familiarization phase and for the four conditions with different probability of action-context co-occurrence in the testing phase. Accuracy values for the familiarization and testing phases were treated, respectively, with between-subjects and mixed model ANOVA designs. Furthermore, in line with previous research (Amoruso et al., 2019; Butti, Corti, et al., 2020), we calculated a standardized beta coefficient across trials of the testing phase by running, at individual level, a regression analysis with probability as predictor and accuracy as dependent variable. This index represents the modulatory effect of the probabilistic associations, thus providing a measure of the strength of the contextual priors. Then, we calculated Pearson’s correlations between the beta index and both the FSIQ and *T*-scores at the social perception subtests within the two clinical groups. Moreover, we used the Fisher’s Z-transformation to test differences between the correlations of the two clinical groups. Finally, we used two-tailed independent samples t-test to analyze differences between groups in FSIQ and social perception abilities.

For Experiment 2, statistical analyses were performed following the same design of Experiment 1 with the addition of task as within-subject variable. Furthermore, a follow-up ANCOVA was used to partial out the effects of IQ. Given the 2 tasks x 4 probabilities x 3 groups design and expecting a medium effect size of f(U)=0.33 based on the results of Experiment 1 (η^2^_p_=0.1), we estimated that a sample size of 54 participants in total ensured appropriate sample after any drop-out (alpha=0.05, power=0.8). Furthermore, for both phases, here we also tested any effect of IQ on the observed group differences in a follow-up Analysis of Covariance (ANCOVA) with IQ as covariate. Statistical analyses of Experiments 1, 2 were performed with Statistica 8.0 (Statsoft, Tulsa, OK). Power analyses were conducted with the G*power software (Faul et al., 2007) using the “as in SPSS” option. We reported ANOVA effect sizes as partial Eta squared (η^2^_p_), adopting conventional cut-offs of η^2^_p_=.01, .06; and .14 for small, medium, and large effect sizes, respectively (Cohen, 2013). Data were reported as Mean ± Standard Error of the Mean (SEM). We set the significance threshold at p=0.05 for all statistical tests. Significant interactions were analyzed with Duncan’s post-hoc test correction for multiple comparisons (Duncan, 1955; McHugh, 2011).

For Experiment 3, we used non-parametric tests for analyses due to small sample size impacting on normal distribution of data. Preliminarily, we verified that the two groups had comparable age, IQ, and gender (Mann-Whitney U/ Chi^2^ tests). For the VR evaluation sessions, we considered the total score and the mean duration of each trial. With the aim to weight the use of predictive and random strategies, we computed, respectively, the mean percentage of scores obtained when the probabilistic avatar-object association gave clues on avatar’s intention (i.e., the 80% avatarobject association trials; prediction score) and the mean percentage of scores obtained when context did not provide reliable information (i.e., the randomly moving avatar and the 10% avatar-object association trials; random score). Data at the action prediction task was extracted as in experiments 1 and 2. Then, we used Mann-Whitney U tests to analyze between-groups differences in performing the action prediction task and the VR evaluation sessions before and after the two trainings. Furthermore, we ran Wilcoxon signed-ranks tests to explore within-group differences. We reported effect sizes using the r index (Z/√N), with the cut-off 0.1, 0.3 and 0.5 for small, medium and large effects, respectively (Fritz et al., 2012). We set the significance threshold at p=0.05 and used the SPSS software for Windows version 21 (IBM corp., New York, NY) for analyses.

## Acknowledgments

The authors are grateful to Elisabetta Ferrari, Daniele Panzeri and Claudio Corbetta for their help in data collection. The authors would also thank all participants and their parents for taking part into this study.

## Data availability statement

The anonymized datasets generated and analyzed during the current study (Experiment 1-2-3) are available from the Open Science Framework, at this link: https://osf.io/wdgsb/?viewonly=92e94c73f9404278bfd9a42031507f5a.

